# Adaptive integration of model-based and model-free strategies in human reinforcement learning of reachable space

**DOI:** 10.64898/2026.03.02.709046

**Authors:** Tianyao Zhu, Rohaan Syan, Shrika Vejandla, Jason P. Gallivan, Daniel M. Wolpert, J. Randall Flanagan

## Abstract

Most skilled behaviour occurs within reachable space; however, how humans learn to reach around obstacles in this space remains almost entirely unexplored. Here, using a novel robotic maze task that captures the richness of naturalistic hand-object interaction, we show that humans adaptively integrate model-based and model-free reinforcement learning strategies to act in reachable space. Fitting hybrid models to reach trajectories revealed that participants shifted from model-based toward model-free strategies across learning. Specifically, model-free reliance increased with state familiarity and distance from the goal, and with exclusive haptic feedback. Across participants, greater model-free reliance was associated with faster movements, consistent with reduced planning demands. Critically, direct comparison with an analogous virtual navigation task revealed stronger model-free reliance in reachable space than in navigable space, demonstrating that the computational architecture governing spatial learning is shared across scales but calibrated to the costs and constraints of the specific effector system.

## Introduction

Many everyday manual tasks—reaching for a coffee cup among dishes, manoeuvring a utensil around obstacles on a cutting board—require selecting and sequencing actions within reachable space, the area immediately surrounding the body where the hands interact with objects [1, 2]. Despite the ubiquity of such behaviour, how representations of reachable space are learned and used to guide manual action remains largely unexplored. Research on reach movement planning and control has focused predominantly on simple environments with limited decision-making demands [3], while the study of spatial learning and decision-making has centred almost exclusively on large-scale navigation. Reachable space thus sits at an unexamined intersection of motor control and spatial cognition.

Navigation research has formalized spatial learning within a reinforcement learning (RL) framework, distinguishing between a model-based (MB) strategy that builds and exploits an internal model of the environment to plan actions, and a model-free (MF) strategy that caches action values through experience [4–10]. MB strategies afford flexible responses to environmental changes but are computationally costly; MF strategies are efficient but inflexible [11–14]. Because neither strategy alone accounts for behaviour, hybrid frameworks combining MB and MF control have been developed [15–22], often modelling behaviour as a weighted mixture of the two strategies [23–25]. The MB/MF distinction is closely tied to the psychological contrast between goal-directed and habitual behaviour [12, 26–31], with the general expectation that MF reliance grows with experience. However, this pattern is not universal: in the widely-used two-step task [24], participants often show the reverse trajectory, relying more on MF strategies early and shifting toward MB strategies later [32]. For reachable space learning, it is unclear how the relative contribution of MB and MF strategies will evolve, as this depends on a complex interplay between cognitive load, motor execution costs, and the inherent uncertainty within both domains. Although dynamic shifts in MB/MF strategies are a central, unresolved question in human and animal RL, hybrid models in practice usually assume a single, fixed weighting, and very few studies implement models that can capture flexible strategy use during learning [33].

There are reasons to expect that the dynamics of MB/MF arbitration in reachable space will differ from navigation. Growing evidence indicates that reachable space is encoded differently from large-scale navigable environments: scene-perception research has identified distinct neural signatures for spaces within reach [34], and dorsal-stream sensorimotor areas preferentially represent objects located within reachable distance [35, 36]. These representational differences likely reflect fundamental asymmetries between the two domains. Navigation relies heavily on visual information to support locomotion over extended spatial and temporal scales, whereas behaviour in reachable space integrates visual, proprioceptive, and haptic information to guide rapid limb movements characterized by trial-by-trial adjustments. The two domains also differ in cost structure: hand movements are biomechanically cheap relative to locomotion, so suboptimal paths carry lower penalties in reachable space. These differences in sensory channels, effector systems, timescales, and movement costs could systematically alter the cost-benefit trade-off governing MB/MF arbitration [27, 31]. The two RL strategies also map onto established motor control constructs, especially regarding limb movements: MB strategies share features with optimal feedback control models [37, 38], which plan actions based on an internal model and ongoing evaluation of outcomes, while MF strategies resemble use-dependent learning [39] and the formation of procedural memories [40].

Here we applied this framework to reachable space using a robotic maze task in which participants moved a handle to reach a target while avoiding obstacles. We compared a Visual–Haptic condition, where participants could see and feel the maze, with a Haptic condition, where the maze was explored solely through touch— contrasting behaviour supported by an immediately available spatial representation with behaviour where a cognitive map must be built through experience. We quantified behaviour using hybrid RL models combining MB and MF algorithms at different levels of granularity and applied the same modelling approach to a previously published spatial navigation dataset [41] to directly compare learning across spatial scales. We show that participants dynamically integrated MB and MF strategies, shifting toward MF control as experience accumulated—a shift modulated step-by-step by state familiarity, distance from the goal, and sensory availability. Direct comparison with the navigation task revealed stronger MF reliance in reachable space, indicating that MB/MF arbitration is calibrated to the biomechanical and informational constraints of the effector system.

## Results

In our reachable space maze task (Fig. 1a, b), participants moved a robotic handle to control a sphere toward a target. The robot simulated contact forces between the sphere and the maze floor, boundary, and blocks. While the target, floor, and boundary were always visible, the maze blocks and hand position were visible only in the Visual-Haptic condition. In the Haptic condition, these elements were hidden, requiring participants to learn the maze layout through haptic feedback.

**Fig. 1.**
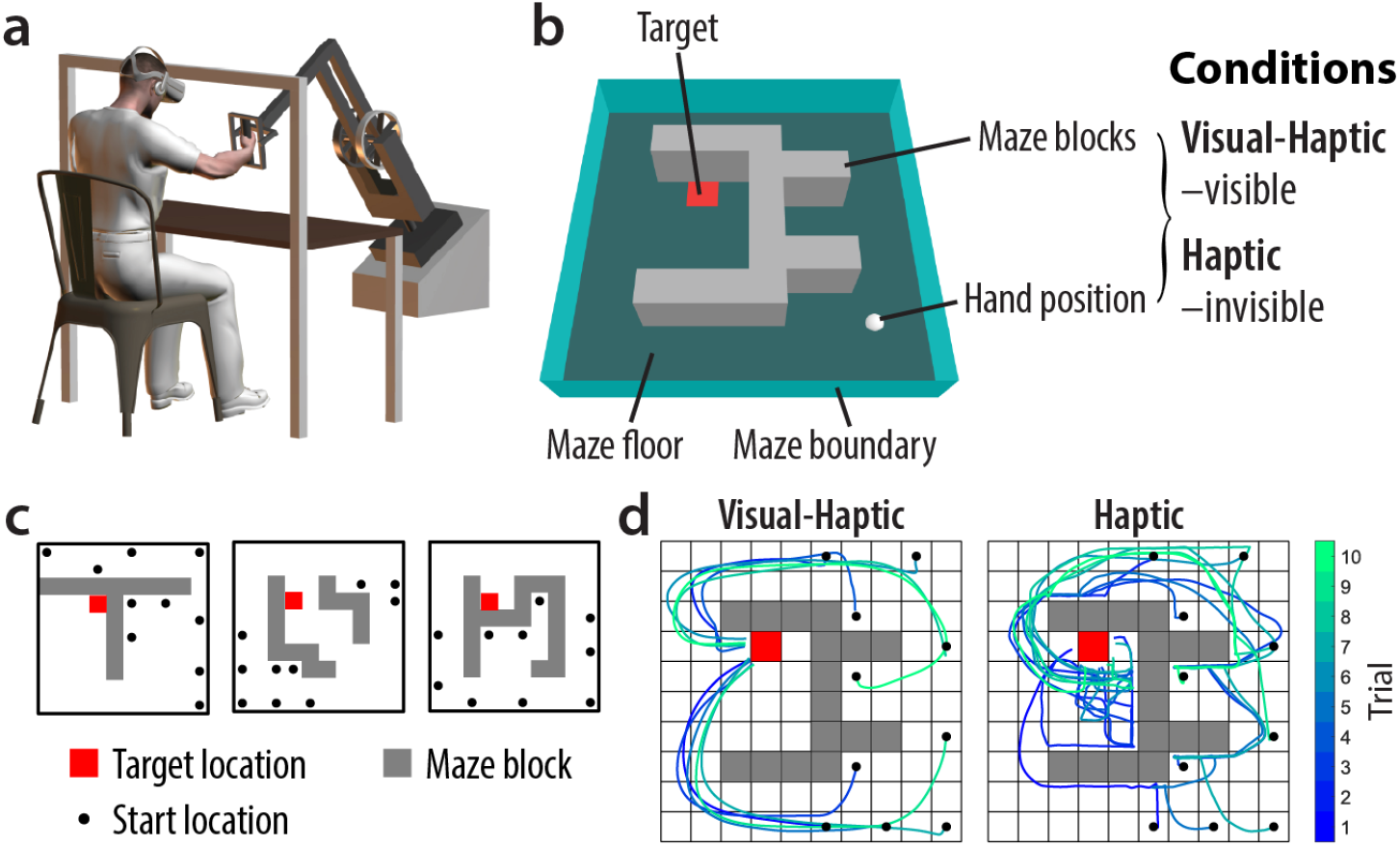
Experimental paradigm and design. a, Experimental apparatus. During the experiment, participants were seated in front of a three-dimensional robotic manipulandum equipped with a virtual reality display. b, Illustration of the virtual maze. In each trial, the participant moved a sphere (white) located at their hand position to a visually cued target (red square), manoeuvring around the maze blocks (grey). Haptic feedback was provided via the robotic handle to the hand when the sphere made contact with the maze blocks, as well as the maze floor (dark green) and boundary (light green). The maze blocks as well as the hand position were visible in the Visual-Haptic condition and invisible in the Haptic condition. c, Example maze configurations. A total of 25 mazes were used per condition of each experiment, with 10 trials performed in each maze. The figure illustrates 3 example mazes. Participants started from different locations (black dots) but always moved to a fixed target location (red square). d, Example participant behaviour. Movement paths are from data of one example participant in each condition. The trajectories were colour-coded by trial number.

Two groups (*N* = 18 each) completed the Visual-Haptic and Haptic conditions, performing 10 trials per maze from various start locations to a fixed target. Figure 1c illustrates 3 of the 25 experimental mazes. Figure 1d shows example movement paths in each condition. In the Visual-Haptic condition, participants typically avoided maze blocks in all trials. In contrast, Haptic condition participants contacted more blocks, especially in earlier trials, as they lacked prior knowledge of the layout at the onset of each new maze.

Prior to each trial, the robot positioned the hand at the start location. Participants then moved the sphere along the floor toward the target, manoeuvring around blocks within a 20 s limit, which was only occasionally reached. Maze blocks and start and target locations were defined in a 10 × 10 gridworld space (see Fig. 1d). See *Methods* for further details on experimental settings and procedures.

### Model-based and model-free reinforcement learning

Raw movement trajectories were discretized into grid paths for modelling. At each *step* along the grid path, the participant took an *action* from their current *state* (one of the 10 × 10 grid locations) to an adjacent state in one of the four cardinal directions (north, east, south, or west). Note that *steps* were defined by transitions between grid locations rather than time-based sampling of trajectories.

We implemented two reinforcement learning (RL) algorithms—model-based (MB) and model-free (MF)—for this reachable space task. In the MB algorithm, we assume participants learn state transition probabilities, representing their belief about whether an action leads to the intended state (no maze block). These probabilities are assumed to be fully learned via visual information at the onset of each maze in the Visual-Haptic condition, whereas they are acquired gradually through touch in the Haptic condition, corresponding to forming a cognitive map. To select an action, the algorithm performs a *planning* process using an offline value iteration method [7, 42] based on current transition probabilities. This method computes a value (expected reward) for each possible action, and the algorithm then applies Boltzmann exploration to select an action. In contrast, the MF algorithm does not learn state transitions. It is implemented as a Q-learning algorithm with eligibility traces [7, 8], which caches values along action sequences, allowing the target reward to propagate back to previous actions. Action selection involves applying Boltzmann exploration directly to these cached values without planning. See *Methods* for a detailed description of the algorithms.

Figure 2 illustrates the characteristics of these algorithms based example participants. The top row shows the value map for each algorithm at the onset of a new maze, before any trials occurred. In the Visual-Haptic condition, the MB algorithm computed a value map based on the actual maze layout. In the Haptic condition, the initial MB map was equivalent to distance heuristics based on a uniform initial transition probability function (the participant’s belief about the overall proportion of empty grids and blocks without specific knowledge of their locations). In the MF algorithm, all initial values were zero; this pessimistic initialization encourages repeating successful action sequences, capturing habitual learning strategies. The bottom row shows the value map learned after all 10 trials, based either on human behaviour or autonomous simulation (see *Methods*). Note that the MB algorithm computed the value map globally through offline planning, given the learned transition probabilities, whereas the MF algorithm only updates values locally along the paths taken and therefore lacks spatial generalization.

**Fig. 2.**
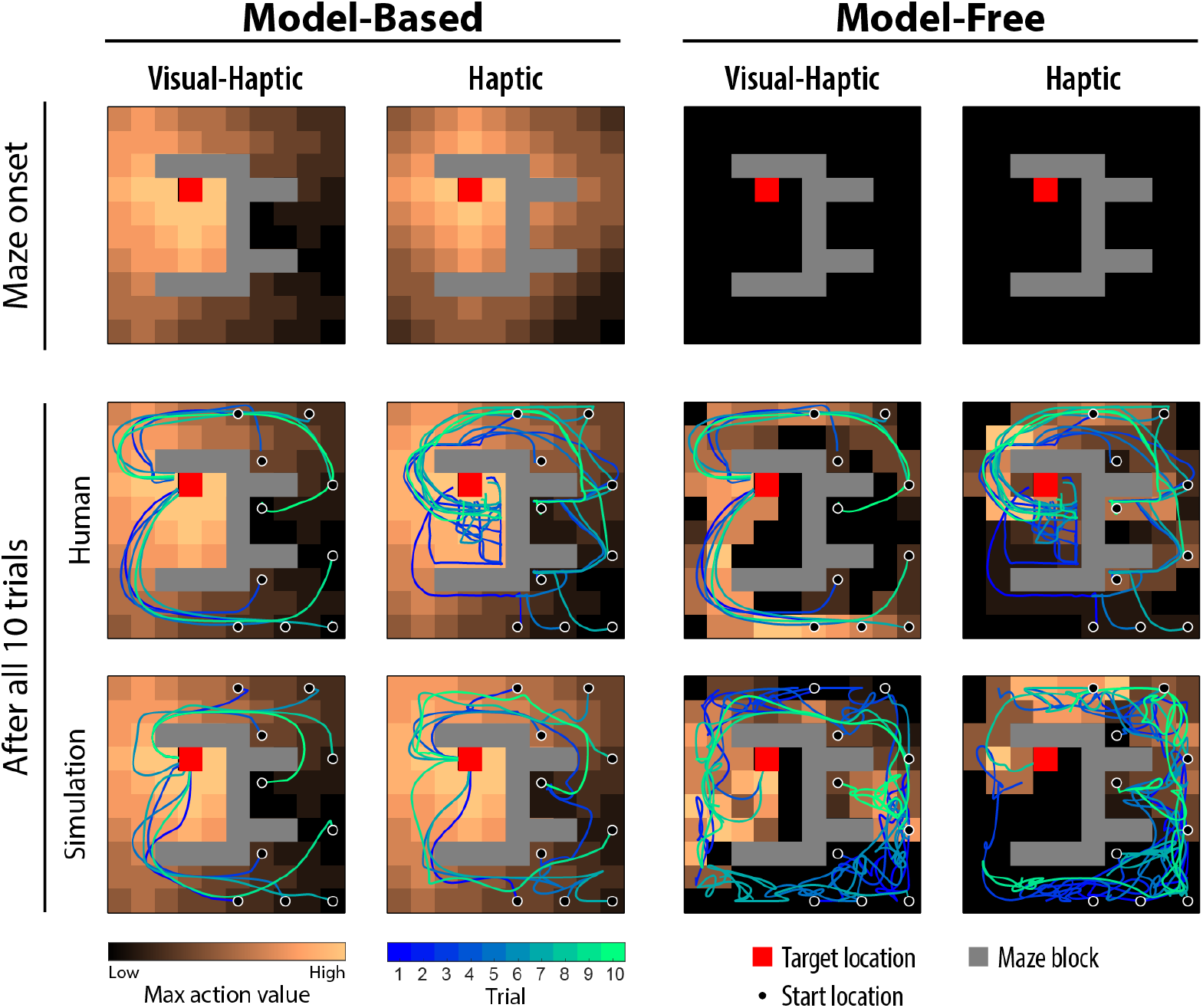
Model-based planning generates global value maps whereas model-free learning updates values locally along experienced paths. Results were generated using the best-fit model parameters for each algorithm based on the example participant’s behavioural data. The colour of each empty, non-target grid represents the highest action value among all possible actions in that state. The top row shows the value map for each algorithm at the onset of a new maze, before any trial occurs. The MB algorithm was able to compute a value map based on actual maze layout (Visual-Haptic condition) or based on a uniform initial transition probability function (Haptic condition), while the MF algorithm initialized all values to zero (pessimistic initialization) before any successful experience. The bottom row shows the value map learned by each algorithm, either based on human behaviour or the autonomous simulation. Simulated grid paths were smoothed to mimic human trajectories. The MB algorithm computed the value map globally through the offline planning process given the learned transition probabilities, while the MF algorithm only updated the values locally along the paths taken. When simulated independent of human data, MB was able to successfully reach the target in most of the trials, while MF failed many times.

### Action strategies in the reachable space maze task

We examined participants’ MB/MF strategies in the reachable space task by modelling their behavioural data (discretized paths). We first fit the MB and MF algorithms separately and compared them to reveal which strategy participants primarily relied on. To address the possibility of a strategy mixture, we also fit a hybrid model using a constant weight to combine MB and MF policies (action probabilities) at each step (hybrid-constant, HC). We further explored whether and how the relative contributions varied during the task. We tested the hypothesis that MB/MF contributions changed monotonically across trials using a hybrid model where the MF weight is a logistic function of trial number (hybrid-dynamic, HD). Moreover, to reveal potential fine-grained changes, we fit a third hybrid model allowing an independent MF weight per action step (hybrid-stepwise, HS). See *Methods* for model details.

Models were fit individually to compute the Bayesian Information Criterion (BIC) [43]. Note that BIC was not computed for the HS model, as it is non-parametric regarding MF weights and unsuitable for BIC evaluation. Figure 3a, i shows the overall BIC (summed across participants) for each model by condition. In the Visual-Haptic condition, the MB algorithm is favoured over the MF algorithm (*BIC*_MB_ = 89, 642, *BIC*_MF_ = 111, 361), while in the Haptic condition, the MF algorithm is favoured (*BIC*_MB_ = 273, 613, *BIC*_MF_ = 267, 105). In both conditions, the HC model is favoured over single algorithms (Visual-Haptic: *BIC*_HC_ = 83, 075; Haptic: *BIC*_HC_ = 254, 270), and the HD model is further favoured over the HC model (Visual-Haptic: *BIC*_HD_ = 82, 112; Haptic: *BIC*_HD_ = 254, 228). The mean logit-slope of the MF weights in the HD model is significantly greater than zero in both conditions (two-tailed one-sample t-test against zero; Visual-Haptic: *M* = 0.21, 95%*CI* = [0.19, 0.23], *p* = 10^*−*14^; Haptic: *M* = 0.058, 95%*CI* = [0.041, 0.074], *p* = 10^*−*6^), indicating a reliable group-level increase of MF weight across trials. This shift from MB to MF strategies during learning occurred in both conditions, including the Visual-Haptic condition where the maze layout was fully visible from the outset—suggesting that the transition reflects a more general reallocation of control as experience accumulates, independent of the reliability of the cognitive map.

**Fig. 3.**
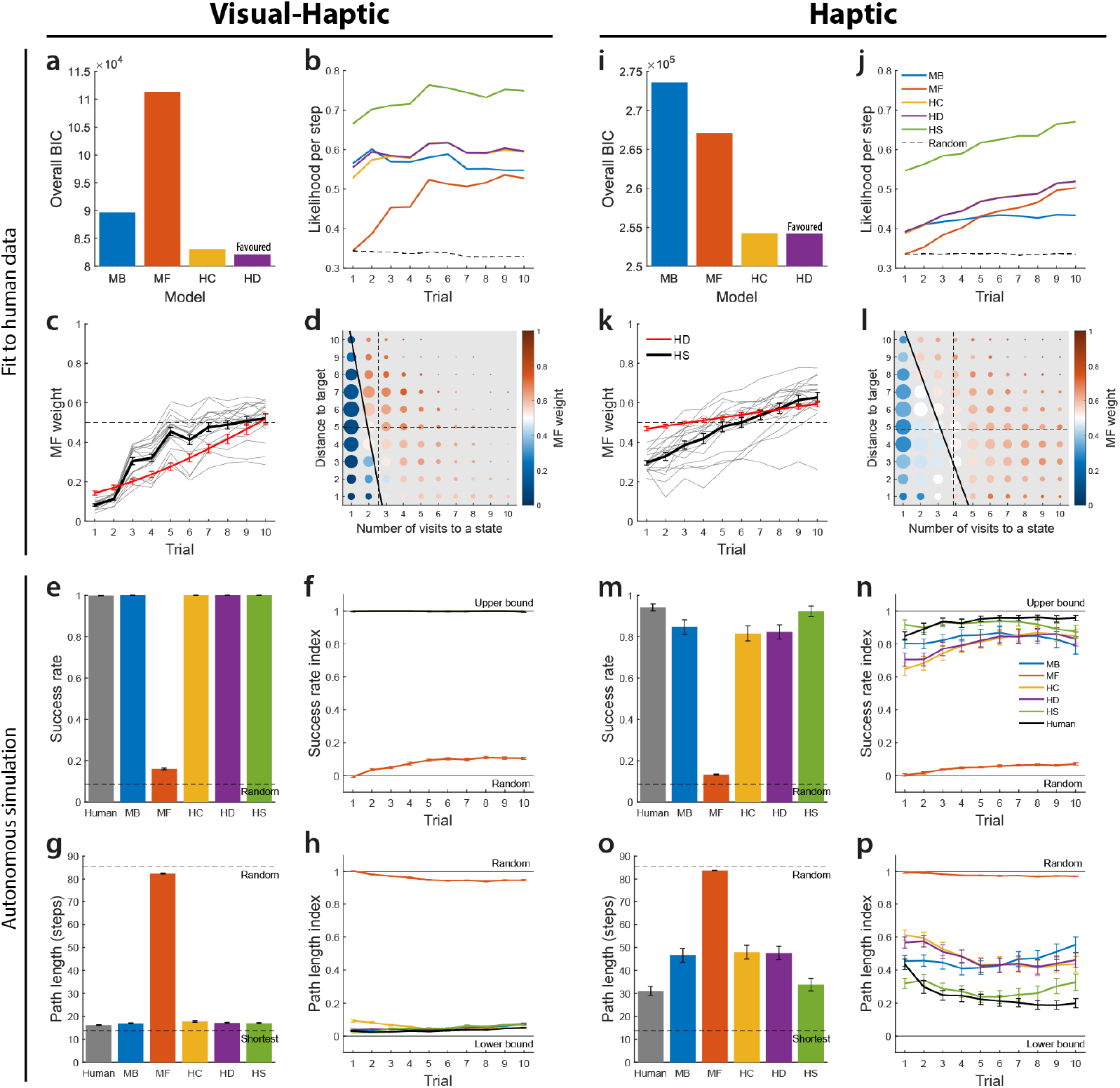
Participants dynamically shift from model-based to model-free strategies during reachable-space learning. a–h, Results for the Visual-Haptic condition. i–p, Results for the Haptic condition. a, i, Overall BIC for each model. Hybrid models (HC, HD) are favoured over single algorithms, with the HD model being the most favoured across both conditions. b, j, Likelihood per step as a function of trial. The MB likelihood remains stable, while the MF likelihood increases across trials. Hybrid models consistently show higher likelihood than single models. c, k, MF weight changes across trials. Average MF weight significantly increased with trial number in both conditions. Overall MF weights were higher in the Haptic condition. Grey lines represent individual participants for the HS model. d, l, Step-wise MF weights plotted against number of visits to a state and distance to the target. MF weight significantly increased with both state familiarity and distance to the target. Dot colour shows the mean MF weight, while dot size is proportional to the number of observations. Dashed lines show the mean number of visits to each state and mean distance to the target across all steps and participants. Thick black line shows a linear approximation of the boundary where the MF weight equals 0.5. e, m, Overall success rate for humans and autonomous model simulations. The single MB algorithm and hybrid models achieve near-human performance, while the single MF algorithm performs near random levels. f, n, Success rate index across trials, scaled between random performance (0) and the upper bound (1). g, o, Overall path length (number of steps) for humans and model simulations. Hybrid models and the MB algorithm match human optimality, whereas the single MF algorithm results in significantly longer paths. h, p, Path length index across trials, scaled between the shortest possible path (0) and random level (1). Error bars show *SEM*.

Next, we examined the average likelihood per action step across trials. Figure 3b, j shows the likelihood per step (averaged across mazes and participants) as a function of trial, using a *random* algorithm as a baseline. The MB algorithm likelihood remained above random and relatively stable, while the MF likelihood started near random in trial 1—as the MF algorithm requires reaching the target to learn effectively—and increased across trials. In the Visual-Haptic condition, the MF likelihood remained lower than the MB likelihood through trial 10, whereas in the Haptic condition, the MF likelihood surpassed the MB likelihood by trial 5. Both hybrid models showed higher likelihood than single models across most trials. The HS model had the highest likelihood, provided for reference as it is not directly comparable to the other models due to its large latent variable vector.

We analyzed the MF weights estimated by the HS model to study task-related strategy changes. We first examined MF weight changes across trials, averaged across mazes and participants (Fig. 3c, k). Using mixed-effects beta regression with a logit link and random intercepts, we found the average MF weight increased with trial number in both conditions (fixed-effect: *p* < 10^*−*16^ for both), indicating a general shift from MB in early learning to MF in late learning. We overlaid the mean MF weight across participants, computed from the parametric logistic function in the HD model, for reference. While the mean MF weights from the HD and HS models are qualitatively similar, the HS model captures more complex structures, whereas the parameterized HD model provides standard statistical inferences regarding the across-trial trend.

Consistent with likelihood results, the overall MF weight estimated by the HS model was significantly higher in the Haptic condition than in the Visual-Haptic condition (two-tailed unpaired t-test, *p* = 10^*−*4^), showing that participants relied more on MF strategies when visual information was unavailable. This is consistent with an uncertainty-based account in which greater ambiguity in the environmental model shifts the balance away from MB planning. Using overall MF weights from either the HC or HD models yielded similar results for this between-condition difference.

To assess control strategies beyond the trial level, we examined how step-wise MF weights depended on two factors: the number of visits to the current state (state familiarity, relevant for MF) and the current Manhattan distance to the target (a proxy for planning complexity, relevant for MB). Heatmaps (Fig. 3d, l) show the average MF weight trends along these factors. Statistical quantification via multiple mixed-effects beta regression confirmed both factors were significant in both conditions (fixed-effect: *p* < 10^*−*16^ for both), indicating participants relied more on MF strategies for familiar states and states farther from the goal. These step-level dynamics suggest that the MB/MF balance is not set globally but is adjusted on a moment-to-moment basis according to local state properties—a degree of granularity not captured by standard hybrid models with fixed or trial-level weightings. Multicollinearity was minimal (Visual-Haptic: *V IF* = 1.11, Haptic: *V IF* = 1.05). The influence of familiarity likely reflects the accumulation of informative MF values, while the distance effect may stem from increased uncertainty in long-distance MB planning [27, 31].

In the analyses above, the models computed values and selected actions conditional on participants’ empirical actions prior to the current step. To examine model performance independent of human data, we ran autonomous simulations for each model using the parameters fit to participants (see *Methods*). Unlike the *fitting* process, during *simulation*, models computed values based on action choices generated by the algorithms themselves rather than empirical actions.

Model performance was evaluated by success rate and path length. For human participants, the success rate is the proportion of trials where the target was reached within 20 s, while path length is the total number of discretized grid steps per trial. For model simulations, we set a maximum step limit per simulated trial based on the typical steps humans took within 20 s. Performance was averaged over 10 independent simulated experiments for each participant. Figure 3e, g, m, o shows mean success rate and path length across all mazes and trials, averaged across participants, for both conditions. We also plotted scaled indices for success rate and path length as a function of trial number (Fig. 3f, h, n, p). Success rate was scaled between 0 (random algorithm performance) and 1 (100% success). Path length was scaled between 0 (shortest possible length) and 1 (expected random algorithm length), facilitating comparisons across trials with different start or target locations.

Overall, the single MB algorithm and all hybrid models performed well in simulations. In the Visual-Haptic condition, these models achieved near-optimal performance, similar to humans. In the Haptic condition, the HS model performance was superior and most closely matched human data, likely due to its fine-tuned MF weights and MB/MF parameters within its flexible hybrid structure. Conversely, the single MF algorithm performed barely better than random and seldom reached the target within the allowed steps in either condition (see Fig. 2 for example paths). This suggests that the MF algorithm, starting from unguided exploration, cannot solve this maze task independently. Notably, although hybrid models relied considerably on MF action choices, particularly in later trials, they performed significantly better than the single MF algorithm. This performance advantage indicates that within the hybrid framework, the MF algorithm updates its values based on successful paths initially generated by the MB algorithm, allowing the MF component to effectively “imitate” these MB-generated actions later. Rather than operating as independent competitors, the two strategies appear to interact cooperatively, with early MB planning scaffolding the MF values that later support efficient, low-cost action selection.

### Individual differences in action strategies and behavioural metrics

MB and MF algorithms typically differ in learning and decision-making behaviour. The MB algorithm generally achieves higher performance, especially during early learning in novel environments, due to its ability to plan flexibly using an environmental model. However, the MB planning process is more computationally costly, often leading to slower responses. Furthermore, the MB algorithm selects actions based on predicted values even without direct experience, naturally exhibiting more “exploratory” behaviour, whereas the MF algorithm repeats previously successful actions, making it more “exploitative.” We therefore investigated whether action strategies could account for individual differences in behavioural metrics like optimality and efficiency.

For each participant, we characterized hybrid strategies using the overall MF weight. Reported results are based on MF weights from the HS model as they are more data-driven, though weights from HC or HD models yield similar results. We assessed optimality via path length (number of steps) and efficiency via average time per step. Importantly, we only analyzed steps without maze block contact (“no-block-contact” steps), as collisions could increase step duration due to biomechanical factors. Metrics were averaged across trials and mazes for each participant.

In the Visual-Haptic condition, where the visible layout supported an accurate mental map for MB planning, path length correlated positively with MF weight (Fig. 4a). Because MB planning produces optimal paths with an accurate map, participants relying more on MF selection produced less optimal solutions. The proportion of no-block-contact steps also correlated positively with MF weight (Fig. 4b). This suggests that MF-reliant participants, by repeating previous movements, may reduce motor variability and accidental collisions. Additionally, average time per no-block-contact step correlated negatively with MF weight (Fig. 4c), indicating faster movement in open spaces. These findings support the idea that MB planning—rather than model learning—is computationally demanding, as the layout was already visible. In the Haptic condition, where the layout was learned through touch, the overall MF weight did not significantly correlate with path length (Fig. 4d). This suggests heavy reliance on the MF algorithm did not compromise optimality, likely because MF selection was primarily used during late learning for states where values were already informative. Path length was significantly longer than in the Visual-Haptic condition (*p* = 10^*−*8^; Fig. 4g), reflecting the lack of an accurate map.

**Fig. 4.**
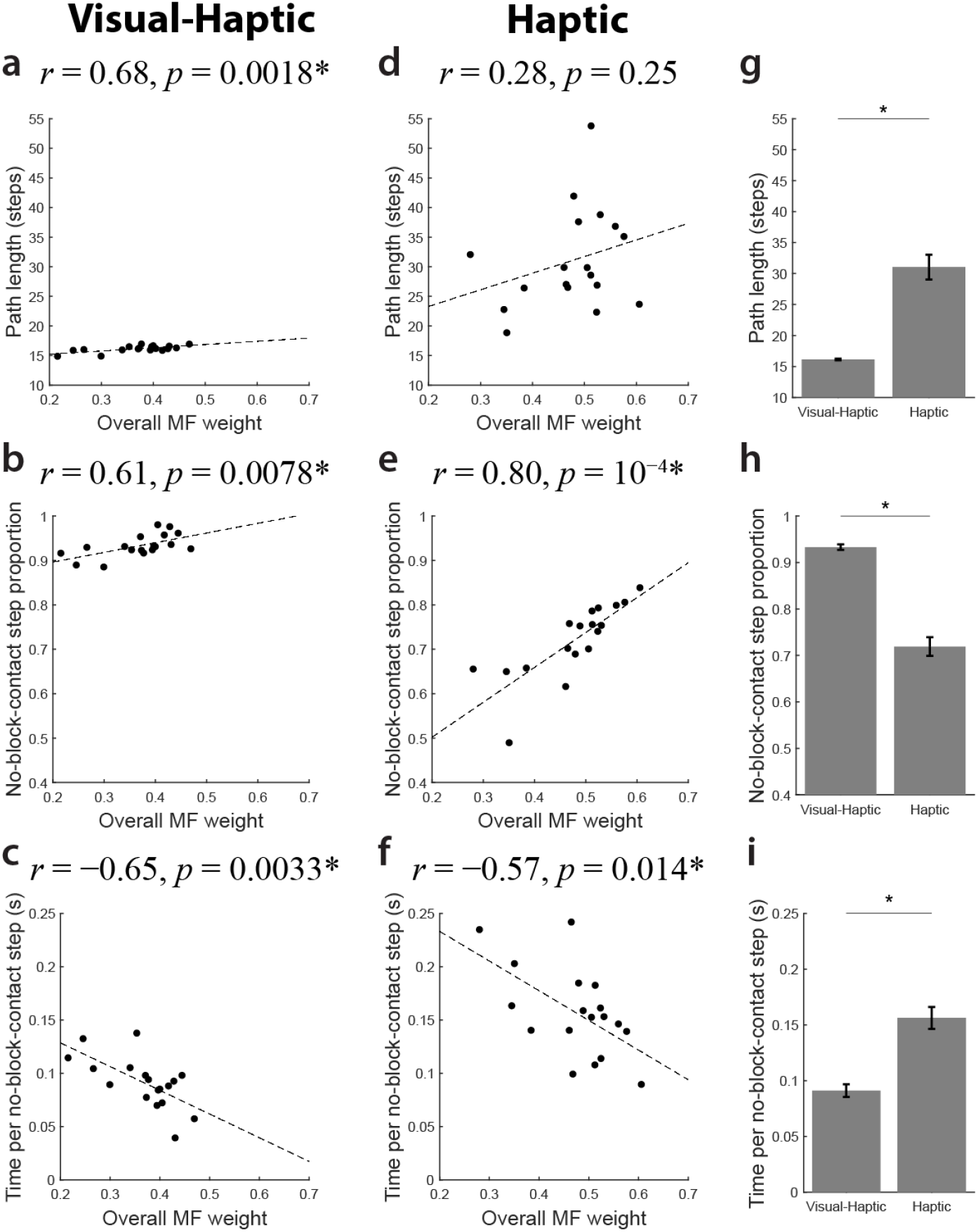
Greater model-free reliance is associated with faster movements and fewer obstacle contacts. a–c, Visual-Haptic condition. Correlations between overall MF weights and a, path length, b, proportion of no-block-contact steps, and c, average time per no-block-contact step. In this condition, higher reliance on MB planning led to shorter, more optimal paths, while MF-reliant participants moved faster and had fewer contacts. d–f, Haptic condition. d, Unlike the Visual-Haptic condition, path length did not significantly correlate with MF weight. Consistent with the Visual-Haptic condition, MF weight correlated positively with e, the proportion of no-block-contact steps and negatively with f, movement time. g–i, Between-condition comparisons. Mean differences between the Visual-Haptic and Haptic conditions for g, path length, h, no-block-contact step proportion, and i, time per step. Participants took longer paths, had more frequent maze block contacts, and moved slower when limited to haptic feedback. Each dot denotes data from an individual participant. Dashed lines indicate linear regression fits. *∗p <* 0.05. Error bars represent *SEM*.

Consistent with the Visual-Haptic condition, the proportion of no-block-contact steps correlated positively with MF weight in the Haptic condition (Fig. 4e). Beyond motor variability, the MF algorithm’s tendency to repeat successful sequences favours open spaces, whereas the MB algorithm’s mental-simulation-based choices may lead to unexpected collisions. Due to lacking visual feedback, the proportion of no-block-contact steps was significantly higher than in the Visual-Haptic condition (*p* = 10^*−*11^; Fig. 4h).

Finally, we again observed a significant correlation between average time per no-block-contact step and MF weight in the Haptic condition (Fig. 4f). Although participants generally favoured MF strategies more than in the Visual-Haptic condition, they moved significantly slower (*p* = 10^*−*6^; Fig. 4i), likely reflecting increased hesitation or caution without visual feedback. An ANCOVA revealed similar slopes across conditions but a higher intercept in the Haptic condition (slope difference: *p* = 0.66; intercept difference: *p* = 0.038), indicating a baseline speed decrease independent of RL strategy use. Collectively, these results suggest that employing MF strategies in the Haptic condition improved efficiency without substantially compromising optimality. More broadly, the association between MF reliance and movement speed across both conditions provides behavioural evidence that the two strategies differ in their real-time computational demands, with MF control enabling faster execution by reducing the need for deliberative planning.

### Action strategies for the navigation task

We applied our modelling approach to data from de Cothi et al.’s virtual reality (VR) navigation task [41] and compared those action strategies to our reachable space results. In the navigation task, 18 participants used a swivel chair and VR controller to move through a virtual maze from various start locations to a fixed target. The maze layout, sequence, start locations, and target were identical to our reachable space maze task. However, instead of blocks rising from the floor, the floor dropped away at those locations (Fig. 5a, left). Only the 3 × 3 grid area surrounding the participant was visible; farther grids were obscured by “fog.” A black curtain marked one boundary to aid orientation. A gold star appeared above the target upon arrival (Fig. 5a, right). Computationally, this navigation task matches our Haptic condition: in both, participants had to move near a grid to obtain information (via limited vision or touch).

**Fig. 5.**
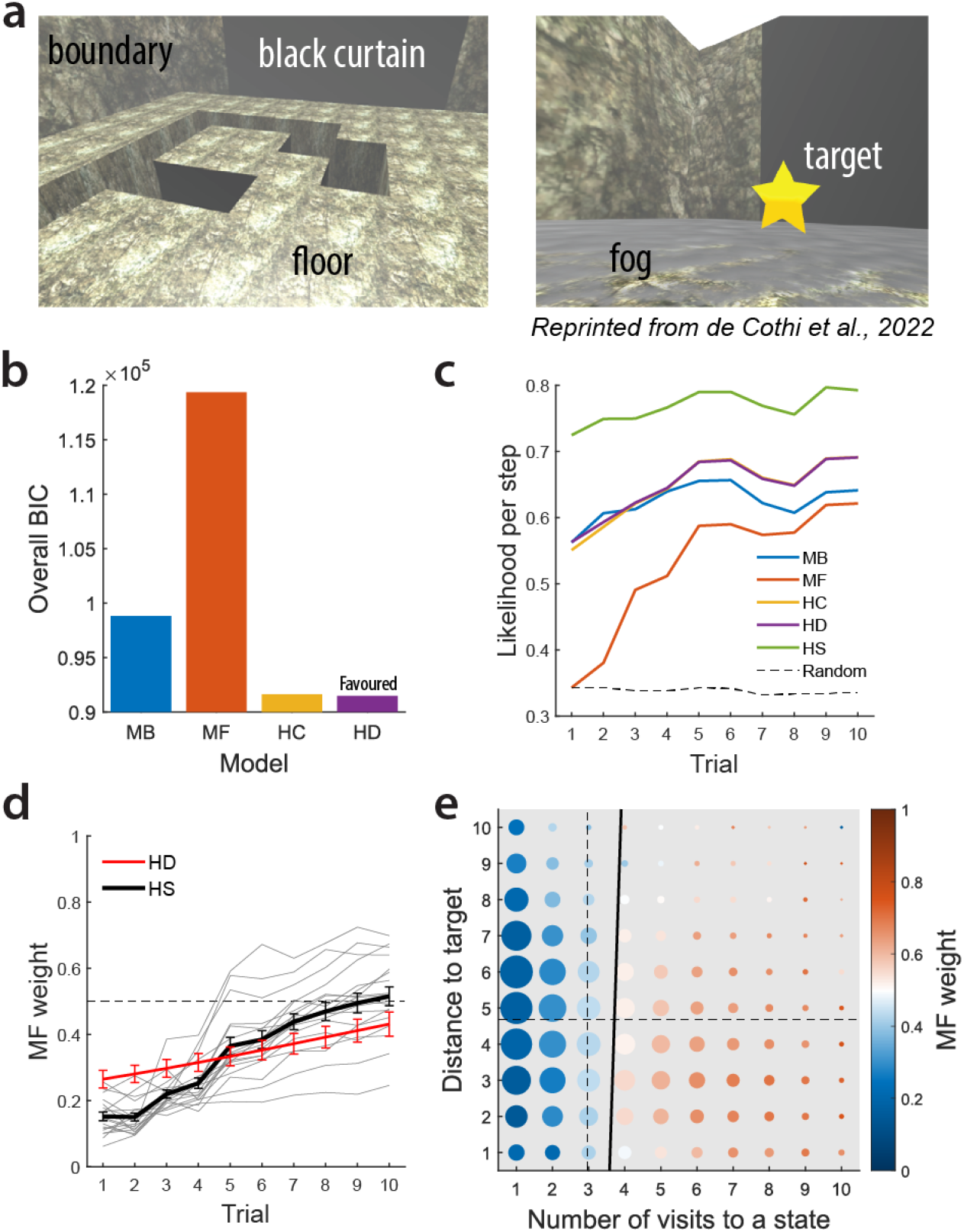
Model-free reliance is weaker in navigation than in reachable space despite matched maze configurations. a, Navigation task environment. Left: the virtual maze environment, including a floor that drops in some locations and a black curtain for orientation. Right: egocentric view in the task. Farther view of the floor was occluded by fog. A gold star appeared above the target upon arrival. b, Overall BIC for each model. Hybrid models (HC, HD) are favoured over single algorithms, with the HD model being the most favoured. c, Likelihood per step as a function of trial. Hybrid models generally show higher likelihood than single models across learning. d, MF weight changes across trials. Average MF weight significantly increased with trial number. Grey lines represent individual participants for the HS model. Error bars show *SEM*. e, Step-wise MF weights plotted against number of visits to a state and distance to the target. MF weight significantly increased with the number of visits to a state, but not with the distance to the target. Dot colour shows the mean MF weight, while dot size is proportional to the number of observations. Dashed lines show the mean number of visits to each state and mean distance to the target across all steps and participants. Thick black line shows a linear approximation of the boundary where the MF weight equals 0.5.

We first examined model comparison results for the navigation task. Figure 5b shows the overall BIC for each model. The MB algorithm is favoured over the MF algorithm (*BIC*_MB_ = 98, 808, *BIC*_MF_ = 119, 355). The HC model is favoured over single algorithms (*BIC*_HC_ = 91, 607), and the HD model is further favoured over the HC model (*BIC*_HD_ = 91, 484). The mean logit-slope of the MF weights in the HD model is significantly above zero (two-tailed one-sample t-test; *M* = 0.090, 95%*CI* = [0.053, 0.13], *p* = 10^*−*4^), indicating a reliable group-level increase of MF weight across trials.

Figure 5c shows likelihood per step (averaged across mazes and participants) as a function of trial. The MF likelihood remained lower than the MB likelihood through trial 10, while hybrid models generally showed higher likelihood than single models.

Analysis of the MF weights from the HS model revealed task-related strategy changes. Figure 5d displays MF weight changes across trials. Similar to the reachable space tasks, the average MF weight increased with trial number (mixed-effects beta regression, fixed-effect: *p* < 10^*−*16^), indicating a shift from MB to MF strategies during learning. However, the overall MF weight estimated by the HS model was significantly higher in our reachable space Haptic condition than in the navigation task (two-tailed unpaired t-test, *p* = 10^*−*5^), showing that participants relied more on MF strategies in reachable space than in navigable space. This cross-domain difference emerged despite identical maze configurations and matched learning structure, pointing to the effector system and its associated movement costs as a determinant of how MB and MF strategies are weighted. The HC and HD models yielded similar results.

In the navigation task, step-wise MF weight significantly increased with the number of visits to a state, but not with the distance to the target (multiple mixed-effects beta regression; fixed-effect for number of visits: *p* < 10^*−*16^, for distance to target: *p* = 0.18; Fig. 5e). Multicollinearity was minimal (*V IF* = 1.07).

### Alternative modelling frameworks

Recent work on human RL [12, 13, 41] has considered successor representation (SR) algorithms, which occupy the spectrum between MF and MB algorithms. SR algorithms learn a predictive map of future state occupancies—the probability that each state will be visited given the current state—which is integrated with reward information to guide action selection [44]. The predictive map can be updated either through temporal difference (TD) learning based on experience (SR-TD), or based on a separately learned transition model (SR-MB) [13]. We evaluated both SR algorithms in our reachable space maze task (see *Supplementary Information*, Fig. S4). In both the Visual-Haptic and Haptic conditions, BIC-based model comparisons show that neither SR model is favoured over the two hybrid models, HC and HD. Specifically, both SR algorithms exhibit lower likelihood than the three hybrid models (HC, HD, and HS) across all 10 trials. These results suggest that SR algorithms are less capable than our hybrid models in explaining participants’ behaviour in these tasks.

In our hybrid models, we used a standard MF algorithm, Q-learning [8], to capture caching or habitual behaviour. However, such behaviour may also be modelled as Hebbian learning [45–47]—an algorithm that strengthens associations based on action frequency—especially when used to “imitate” the actions of a more flexible learning system [48]. We opted not to separately test an MB-Hebbian hybrid because, in this imitation context, it would not offer significant behavioural divergence or additional explanatory power. Furthermore, maintaining the Q-learning framework ensures better compatibility with standard RL benchmarks.

## Discussion

Here we demonstrate that humans adaptively integrate model-based (MB) and model-free (MF) reinforcement learning (RL) strategies to learn reachable space—a domain of spatial cognition that, despite underpinning most everyday skilled behaviour, has received almost no computational investigation. Using a novel robotic maze task and a hybrid modelling framework that tracks the dynamic MB/MF contribution, we show that participants shifted from MB toward MF control across learning trials. This shift occurred not only when participants had to build a cognitive map through haptic exploration (Haptic condition), but also when the maze layout was immediately and fully visible (Visual-Haptic condition)—indicating that the transition to MF control is driven by the intrinsic cost of MB planning, not solely by uncertainty in the environmental model. Whereas RL-based accounts of spatial learning have been developed almost exclusively in the context of navigation [4, 5, 10, 18, 19, 21], we further show through direct comparison with an analogous navigation task [41] that the balance between MB and MF control differs quantitatively across spatial domains, with stronger MF reliance in reachable space. Together, these findings establish reachable space as a tractable domain for studying RL-based action selection and indicate that the computational logic of spatial learning varies with the scale, sensorimotor modality, and cost structure of the behaviour it supports.

Beyond the general MB-to-MF strategy shift across time, we also observed other factors that influence the strategy balance. We found that participants relied more on 16 MF strategies in the Haptic condition compared to the Visual-Haptic condition. Stepwise analyses further revealed that MF reliance was higher for more familiar states and for states farther from the goal. These findings can be unified under the notion of uncertainty-based integration of MB and MF strategies [27, 31, 49, 50], i.e., strategies with lower uncertainty are favoured. Specifically, lacking visual information and planning over longer distance increases the uncertainty in MB strategies, while lacking experience increases the uncertainty in MF strategies. This adaptive arbitration enables participants to maintain high task performance while reducing computational costs when possible. Whether a “meta-controller” governs this arbitration by evaluating the relative reliability of each strategy [30, 51–53] or whether the two systems compete directly [54–56] remains an open question.

While this descriptive account of strategy use is inferred through computational modelling, the MB and MF algorithms map onto cognitive and motor processes in interpretable ways. The hallmark of MB strategies is the capacity to select actions via planning based on an internal model of the environment, enabling flexible generalization across states and actions beyond directly reinforced experience [27, 28, 57–59]. In reachable space, this corresponds to moving the hand toward a target from a novel location based on the seen or remembered spatial layout—a process that becomes slower and less accurate as the number of steps to plan increases [10, 17, 31, 60, 61]. The interpretation of MF strategies is more nuanced. MF control can reflect a spectrum ranging from caching action values and using them to guide future choices [24, 62], to automatic, habitual behaviour that is no longer sensitive to value [39, 47, 63–65]. Along this spectrum, response time decreases, flexibility reduces, and the underlying process shifts from explicit to implicit [66–68]. Computational modelling alone usually cannot distinguish between these interpretations, as they yield similar behavioural predictions; future work combining selective task manipulations with neural evidence may help dissociate them.

Importantly, we found that participants who relied more on MF strategies generally moved faster through the maze. This behavioural correlation with model-estimated parameters supports our interpretation of the MB and MF algorithms as the slow, deliberate planning-to-goal process and the fast, automatic stimulus–response process, respectively. Another intriguing behavioural association is that participants with higher MF reliance tended to contact fewer maze blocks as they moved through the maze. This finding points to an additional benefit of MF control in motor tasks: MB strategies select actions based on simulated values and can route the hand through unexplored regions where the cognitive map is inaccurate, leading to unexpected collisions. MF strategies, by contrast, favour repetition of previously successful actions and produce more conservative trajectories. Movement repetition may also reduce variability in limb motor execution through practice [69–71], further decreasing accidental contacts.

Model simulations provided additional insight into how MB and MF strategies work together in practice. We found that the MF algorithm alone rarely reached the target, indicating that early in learning, participants relied primarily on MB planning to generate successful paths. As experience accumulated, the MF algorithm became increasingly viable by effectively caching the action sequences that MB planning had initially produced. This pattern—MF “imitating” MB-generated solutions—is consistent with the broader proposal that habits crystallize around actions originally selected through goal-directed planning [12, 26–31]. The finding that greater MF reliance predicted faster movements further suggests that the brain does not compute full MB plans on every step but selectively engages planning when its expected benefit outweighs its cost [16, 60]. That this pattern held even in the Visual-Haptic condition—where the environmental model was accurate from the onset—reinforces the interpretation that planning cost, in addition to model uncertainty, is a primary driver of the shift toward MF control. When MF control dominates, planning is curtailed, freeing computational resources and speeding responses. At a broader level, the MB-to-MF transition we observe formalizes, within an RL framework, a progression that motor learning research has long described qualitatively—the shift from slow, deliberative control to fast, automatic performance as a skill is acquired [72, 73].

By comparing the haptic version of our reach space task—without vision of the maze—to an analogous navigation task [41], we could directly compare reinforcement learning strategies in reachable and navigable space. Strikingly, we found that participants relied more heavily on MF strategies in the reachable space task compared to navigable space task, despite the two tasks sharing identical maze configurations and learning structure. We attribute this difference to the distinct cost-benefit landscape of manual action. Hand movements are much faster than locomotor movements: each additional step in reachable space incurs a smaller time cost, reducing the marginal benefit of extensive MB planning to optimize the path. Proprioceptive and haptic feedback during reaching may also facilitate the encoding and reproduction of successful state-action sequences, accelerating the accumulation of informative MF values. These observations point to a broader principle: MB/MF arbitration is not a fixed property of the learner but is shaped by the biomechanical and informational constraints of the effector system through which behaviour is expressed.

These findings raise the question of what neural circuits support MB and MF control in reachable space. Neurophysiological work on reach planning has not generally distinguished between these strategies, and it remains unclear whether motor, premotor, and parietal areas are differentially engaged during MB versus MF action selection. A particularly open question is whether reaching movements in rich environments— with multiple objects and alternative movement paths—engage hippocampal circuits. In navigation, MB strategies depend on cognitive maps represented in hippocampal circuits [74–77].Whether analogous maps of reachable space are also encoded in the hippocampus is unknown. Anatomical connections between medial temporal lobe structures and premotor cortex [78–81] could provide a pathway through which map-like representations inform motor planning in reachable space. Alternatively, the internal models supporting MB reach planning could reside in parietal-premotor circuits that maintain representations of limbs and objects near the body [35, 36, 82–84]. Resolving the degree to which these processes recruit sensorimotor versus hippocampal maps represents an important target for future neuroimaging and lesion studies.

Our findings also generate testable predictions for clinical populations. Disorders affecting basal ganglia function and dopaminergic signaling, such as Parkinson’s disease, produce well-documented disruptions in motor learning and RL [85–88], and our results predict that these disruptions should affect learning in reachable space. Similarly, conditions associated with maladaptive MB/MF control in abstract decision tasks, such as obsessive-compulsive disorder [55, 89–91], may be extended to reachable space tasks as well. Conditions affecting spatial cognition, such as temporal lobe epilepsy [92] and stroke [93, 94], may show distinct patterns in reachable space and navigable space tasks. The robotic maze paradigm and computational framework developed here provide a foundation for examining how these conditions affect behaviour in the space that matters most for daily life—the space within arm’s reach.

## Methods

### Participants

We collected data from 18 participants (mean age 20.4 years, range 18–24 years; 11 females) in the Visual-Haptic condition and another 18 participants (mean age 22.4 years, range 18–30 years; 9 females) in the Haptic condition. These sample sizes align with those typically used in studies of human motor and spatial learning and were sufficient to detect the key within- and between-condition effects. See *Supplementary Information*, Fig. S5 for analysis controlling for the effect of sex when examining the effect of condition. Participants reported no history of neurological disorders and had normal or corrected-to-normal vision. All participants were compensated $15 CAD for their participation. Written informed consent was obtained from all participants. The experiment was approved by the Queen’s General Research Ethics Board and complied with the Declaration of Helsinki.

### Apparatus and stimuli

During the experiment, participants were seated in front of a custom three-dimensional robotic manipulandum (3BOT, Fig. 1a) equipped with an integrated virtual reality (VR) head-mounted display (Oculus Rift DK2). The display offered a resolution of 960 *×* 1080 pixels per eye, a refresh rate of 75 Hz, and a response time of 2 ms. Participants used their dominant hand to grasp the manipulandum’s handle, which was driven by DC motors capable of applying forces to simulate the virtual maze. High-resolution encoders tracked the handle’s position at 1000 Hz, while the forces applied to the handle were recorded at the same frequency. The maze was positioned roughly at chest level and centred in front of the participant. The virtual reality space was aligned with the physical robotic workspace, ensuring consistency between the visual and haptic feedback.

The virtual maze (Fig. 1b) consisted of a 10 × 10 grid array (grid width 2 cm), with a floor (dark green) at the bottom and rising boundary (light green). Each grid square could be either reachable (empty space) or unreachable (filled with a maze block, shown in grey, rising from the floor). Participants interacted with the maze environment through a virtual sphere (white) with a 1 cm diameter—located at the robotic handle grasped in their hand. Counterforces were generated by the manipulandum when the sphere made contact with the maze floor, boundary and blocks, providing haptic feedback. The simulated floor also supported the participants’ hands to reduce fatigue. The maze floor and boundary remained visible to the participant throughout the experiment, filled with a uniform colour and with no gridlines displayed. The maze configurations were adopted from a previous rat and human navigation study [41].

### Experimental procedures

The experiment began with a demo session consisting of 2 phases. Phase 1 included a single trial lasting 20 s, during which a visible example maze was presented (similar, but not identical, to the mazes used in the learning sessions). The sphere was also visible, but no target was present in this trial. Participants were instructed to move the sphere around the maze, feeling the floor, boundary, and blocks of the maze to familiarize themselves with the virtual environment. Phase 2 included 10 trials in which there were no maze, but a visible target location (filled red) was present in the workspace. Participants were instructed to move the sphere to the target location. Participants started from a different start position in each trial and moved to the same target position used in the learning sessions. At the beginning of each trial, participants’ hands were lifted above the floor, directed to the start position, and lowered onto the floor by the manipulandum. Participants then moved to the target on their own. When the target was reached, a beep would sound, and there would be haptic feedback simulating sinking through target grid location. The trial would then end, and the participant would be guided to the start position of the next trial by the manipulandum. In the Visual-Haptic condition, the sphere was visible in all 10 trials. In the Haptic condition, however, the sphere was only visible in the first 5 trials and became invisible in the last 5 trials. Phases 2 was designed to help participants gradually familiarize themselves with the trial flow and the target feedback described above. Specifically, the last 5 trials in the Haptic condition enabled participants to practice reaching the target location without visual feedback of their hand position.

After the demo session, participants began the learning session, during which they would learn 25 different mazes. The maze blocks together with the hand position were visible in the Visual-Haptic condition while they were invisible in the Haptic condition. Participants performed 10 learning trials in each maze (Fig. 1c): they moved from 10 different start positions (black dots) to a fixed target position (red square). The target was visible throughout the trial. Trials terminated either when the participant reached the target or when the maximum allowed time of 20 s was reached. Between maze changes, participants’ hands were lifted and moved outside of the maze boundary and then guided back by the manipulandum, signalling that the maze had changed. The mazes, start positions, and their order were identical for all participants. Participants were allowed to take a short break between different mazes if needed.

### Data analysis

The raw movement trajectories of the participants’ hands were discretized into grid paths, and the virtual maze blocks contacted at each step (i.e., transitions between a grid and one of its four orthogonally adjacent grids) were identified.

### Models

We developed individual model-based (MB) and model-free (MF) reinforcement learning (RL) models, as well as different hybrid RL models to account for the participants’ behaviour.

### Model-based algorithm

In the MB learning algorithm, we assume that participants maintain a mental representation of the maze in the form of a transition function *T* (*s, a*), which denotes the probability that by taking action *a* (going north, east, south, or west) from grid state *s*, the intended state *s*^*′*^ is reachable (no maze block in the grid). That is, we assume that participants understand how their current state and action lead to a transition on the 10×10 grid (i.e. to *s*^*′*^) in the absence of any maze. Note that this probabilistic transition function captures participants’ uncertainty about the maze, while the maze itself is deterministic in our task.

In the Visual-Haptic condition, we assume that the visual maze map is used to construct a true transition function *T* (binary encoding for reachable/unreachable intended states). This transition function does not change across trials.

In the Haptic condition, however, the maze layout is invisible and must be learned through haptic feedback, formalized as follows.

At the onset of a new maze, we assume that the initial transition function *T*_0_ takes a uniform value *p*_0_ between 0 and 1 for all state-action pairs that do not lead outside the boundary (and a value of 0 for those that do). A large *p*_0_ value means that the participant believes that there are more spaces and less blocks in the maze, and vice versa.

The transition function is updated each time the participant interacts with an intended state *s*^*′*^ (entering the grid location or contacting a maze block). The algorithm first looks up all possible state-action pairs (*s, a*) that lead to state *s*^*′*^ (four if *s*^*′*^ is not next to the boundary). This reflects a straightforward spatial inference consistent with a cognitive map of the physical space: if a grid is reachable or unreachable from one side, it will be consistently reachable or unreachable from all other sides (ignoring sides that have blocks, which are therefore never entered). The transition function value for each pair is then updated with learning rate *α*:

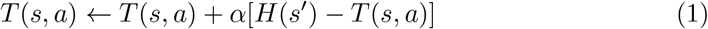

where *H* encodes the sensory information about the intended state:

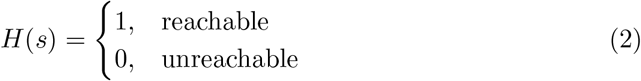

The algorithm separately represents the target location with a reward function *R*:

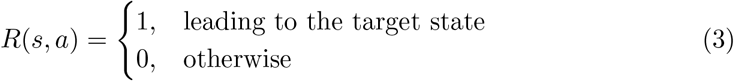

In both the Visual-Haptic and Haptic conditions, to select an action, the algorithm performs planning based on the transition function *T* and the reward function *R*. Specifically, it uses value iteration, a dynamic programming method to approximate the optimal value function by solving the Bellman equation iteratively. In this process, it first initializes the value function *V* arbitrarily (e.g., set to 0) for each state *s*. We set *T* to 0 for all actions originating from the target state (treating it as a terminating state). Then, the following is repeated until convergence:

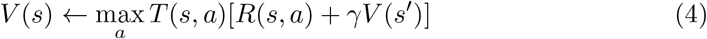

where *s*^*′*^ is the state that (*s, a*) leads to, and *γ* is the temporal reward discount factor. Note that the iterations described above are performed as a separate *offline* (simulated) process, executed before each *online* (actual) step is taken. This differs from the iterative, step-wise learning for the transition function *T* in the MB algorithm and the Q-function in the MF algorithm (see next section).

The probability of selecting each action option *a*_*i*_ from state *s* is:

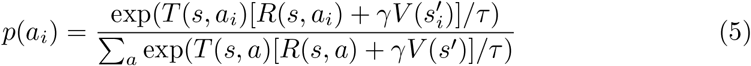

where *s*^*′*^ is the state that (*s, a*) leads to, *s*^*′*^_*i*_ is the state that (*s, a*_*i*_) leads to, and *τ* is the temperature parameter for Boltzmann exploration (softmax). A high temperature indicates more exploration (more random choices), while a low temperature favours exploitation of the best-known actions.

### Model-free algorithm

As the MF algorithm lacks a representation of the transition function, it cannot utilize the visual information of the maze layout in the Visual-Haptic condition. Consequently, the formalization of the MF algorithm does not differ between the two conditions.

Specifically, in this algorithm, we assume that participants maintain a Q-function *Q*(*s, a*), which is the cached value (expected cumulative reward) that can be obtained by taking action *a* at state *s*. At the onset of a new maze, we assume that the initial Q-function *Q*_0_ is 0 for all state-action pairs. With this initial condition, the algorithm is more likely to repeat actions that previously led to rewards as it learns, effectively modelling the formation of habits over time. Because MF learning may represent an automated motor process, it is possible that learned values from previous mazes carry over into new environments. We addressed this by testing two simplified possibilities for the Q-function: a “reset” version (clearing at each new maze) and a “running” version (inheriting values throughout all mazes). The “reset” model provided a better fit for the data in both conditions. This indicates that participants effectively partitioned their learning between mazes, leading us to utilize the “reset” version in our analysis.

The Q-function over all state-action pairs (*s, a*) is updated each time the participant takes an action *a*_*t*_ from state *s*_*t*_ at step *t*:

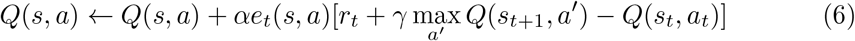

where *r*_*t*_ is the immediate reward received at step *t* (1 if the target state is reached, and 0 otherwise). Note that, unlike MB learning, which represents a reward function *R* to plan actions, MF learning obtains reward information through interaction with the environment, without explicitly representing it in the algorithm. max_*a*_^*′*^ *Q*(*s*_*t*+1_, *a*^*′*^) is the expected future reward, discounted by the factor *γ*. The term *r*_*t*_ + *γ* max_*a*_^*′*^ *Q*(*s*_*t*+1_, *a*^*′*^) − *Q*(*s*_*t*_, *a*_*t*_) is the reward prediction error at step *t*. The Q-function is then updated based on the reward prediction error with learning rate *α* and eligibility trace *e*, which serves as a memory buffer for previous state-action pairs, allowing learning to propagate through the action sequence. The eligibility trace is initialized over all state-action pairs (*s, a*) at the beginning of each trial:

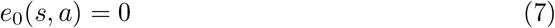

and updated at each step *t* as follows:

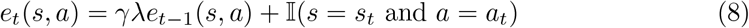

where *γ* is the temporal discount factor for reward, *λ* is the memory decay factor, and 𝕀 is the indicator function, which takes the value of 1 if the condition holds, and 0 otherwise.

During action selection, the algorithm makes decisions based directly on the current Q-function. The probability of selecting each action option *a*_*i*_ from state *s* is:

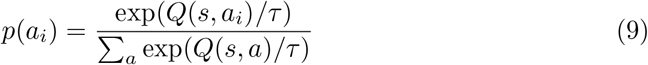

### Parameter fitting

We fit the two algorithms separately for each participant. Both the MB and MF algorithms update their representations at each step as described above based on the behaviours (entering grids and contacting maze blocks) of the participant. For each step the participant entered a new grid, the probability that the agent would select the same action as the participant did (out of all possible actions leading to reachable states) was calculated. We fit the parameters (MB: *γ* and *τ* in the Visual-Haptic condition, and *α, γ, p*_0_, and *τ* in the Haptic condition; MF: *α, γ, λ*, and *τ* in both conditions) to maximize the data likelihood (i.e. minimize the sum of the negative log-likelihood across steps) using *fminsearchbnd* (bound-constrained optimization) in MATLAB [95]. We selected the best-fit parameter set from 10 optimization runs with random initial values. See *Supplementary Information*, Fig. S1a and b for parameter and model recoverability tests for our MB and MF models.

### Hybrid models

In the hybrid RL models, we assume that during the experiment, both the MB and MF algorithms learn in parallel from the experiences generated by the participant, but that the action choice *a*_*t*_ at each step *t* was generated by only one algorithm. Let *z*_*t*_ denote the latent indicator variable specifying which algorithm generated the action.

We define the MF weight for each choice (*w*_*t*_) as the prior probability that the choice was generated by the MF algorithm:

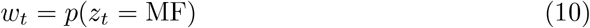

We developed three hybrid models with different assumptions regarding the structure of the MF weight. In the simplest hybrid-constant (HC) model, we assume that the MF weight is constant across all steps:

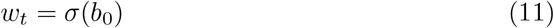

where *σ* denotes the sigmoid (logistic) function:

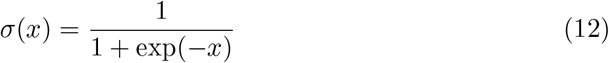

In the hybrid-dynamic (HD) model, we assume that the MF weight is a generalized linear function of the trial number:

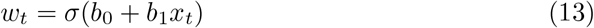

where *x*_*t*_ denotes the ordinal index of the trial (within each maze) to which step *t* belongs.

Lastly, in the hybrid-stepwise (HS) model, the MF weight is not parameterized; instead, each step has an independent MF weight *w*_*t*_.

Given the participant’s choices and the parameters of the two algorithms, *θ*_MB_ and *θ*_MF_, we define the posterior probability (responsibility) of the MF algorithm as:

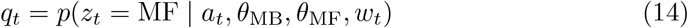

We used Expectation-Maximization (EM) [96] to jointly estimate the MF weights (or their parameters) and the algorithm parameters *θ*_MB_ and *θ*_MF_. In each EM iteration, we computed a lower-bound on the log-likelihood:

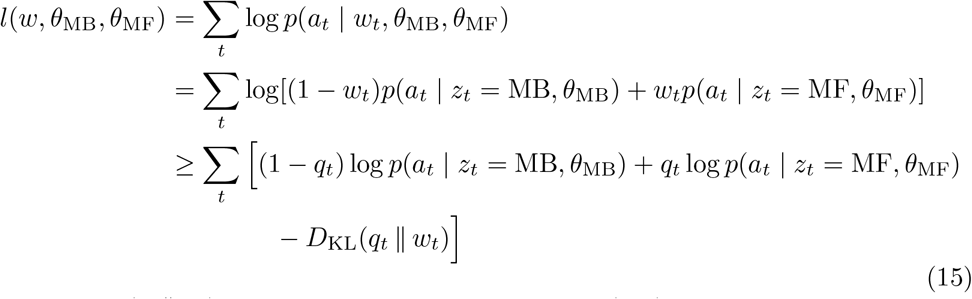

where *D*_KL_(*q*_*t*_ ∥ *w*_*t*_) denotes the Kullback–Leibler (KL) divergence between two Bernoulli distributions with parameters *q*_*t*_ and *w*_*t*_.

We initialized the MF weights *w*_*t*_ to 0.5 for all steps (*b*_0_ = 0 in the HC model and *b*_0_ = *b*_1_ = 0 in the HD model). We also initialized *θ*_MB_ and *θ*_MF_ to the parameters obtained from independently fitted MB and MF algorithms.

In the E-step, we computed the posterior responsibilities of the MF algorithm:

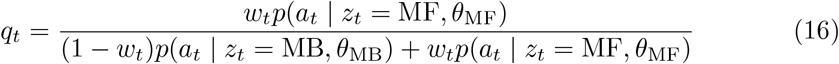

In the M-step, we used maximum likelihood estimation (MLE) to update the parameters of each algorithm separately:

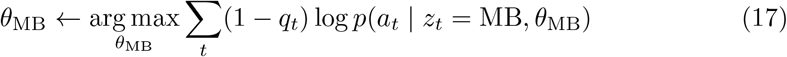

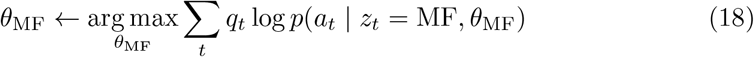

We also updated the MF weights. For the HC model, we updated the constant parameter as:

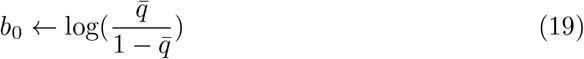

where 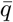 is the mean responsibility across all steps.

For the HD model, we updated the parameters *b*_0_ and *b*_1_ by fitting a logistic regression model to the responsibilities of all steps, using trial number as the predictor.

For the HS model, we set the MF prior weight for each step in the next iteration to the posterior responsibility from the previous iteration,

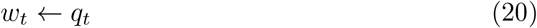

We iterated the E-step and M-step until the relative change in the data log-likelihood was smaller than 10^*−*4^.

See *Supplementary Information*, Fig. S1c and Fig. S2 for validation of the hybrid models.

### Autonomous simulation

In the fitting procedures described above, the model makes predictions of the action, but these predictions were not used. Instead we used the participants’ actual actions when fitting the models.

To examine how the models perform when choosing their own actions, we also simulated the MB, MF and hybrid models in which the algorithm learned by observing outcomes of its own action choices rather than the participants’.

At each step, the algorithm first considered all possible actions that did not lead outside the maze boundary. An action was drawn from the probabilistic distribution over these options calculated by the algorithm. If the action led to a maze block, it was marked as “contacted,” and, in the Haptic condition, the transition function in MB was updated accordingly. The algorithm would then select another action from the remaining options. This process was repeated until the selected action led to a reachable state. The algorithm then “entered” this state, updated its representation, and proceeded to the next step. Each simulated trial ended either when the algorithm arrived at the target state or when the maximum number of steps was reached. Simulated grid paths as well as simulated contacts with maze blocks in each step were recorded.

The maximum number of steps was calculated by averaging the grid path length across all out-of-time trials from all participants (88 steps). This is the approximate number of steps participants could typically take within the time limit of a single trial.

For the separate MB and MF learning algorithms, we ran simulations under each participant’s best-fit parameters. For the hybrid models, the fitted MF weights were used to combine the probabilistic distributions over the options calculated separately by the MB and MF algorithm, then an action was drawn from the combined distribution in each step. Note that the HS model, in which the fitted MF weights were bound to participants’ actual actions, is a descriptive rather than generative model. In order to run autonomous simulations, we used an approximated generative version of the descriptive HS model. Specifically, we used a single average MF weight (i.e., individual data points in Fig. 3c and k) for all steps within each corresponding trial to generate actions.

### Models for the navigation task

In de Cothi et al.’s navigation task [41], participants received visual feedback regarding the maze structure through virtual reality scenes (Fig. 5a). At each step, a 3 ×3 grid array centred on the grid the participant was currently in became visible. In the MB algorithm, we assume that at the first step of each trial, the transition function is updated for all these 9 grids, as the participant would typically need to look around to orient themselves in the maze. At each subsequent step, the transition function is updated for the 3 grids that the participant has newly seen (front-left, front-centre, front-right). Other model settings were similar to those of the reachable space maze task.

## Acknowledgements

We gratefully acknowledge Martin York for programming and hardware support for the experiments. We would also like to thank Dr. Peter Dayan for valuable feedback and suggestions on this work.

## Supplementary Information

### Recoverability tests

We performed parameter recovery for the parameters of the model-based (MB) and model-free (MF) algorithms. Tests were performed separately for the Visual-Haptic and Haptic conditions. For each participant, we ran autonomous simulations with the MB and MF algorithms separately using their best-fit parameters (“true” parameters). Then, we fit the same model (MB or MF) to the simulated data and obtained a set of “recovered” parameters.

Fig. S1a shows the recovered parameters against the true parameters. Each dot represents a single participant. For most model parameters, the recovered parameters were close to the true parameters (the dots are close to the dashed *x* = *y* unity line), with high Pearson’s *r* values, showing good recoverability. The *λ* parameter in the MF algorithm in the Haptic condition, which is the memory decay parameter for eligibility traces, had a relatively low *r* value. Note that this parameter was close to 1 for all participants, which made it difficult to recover the small between-participant differences. Nevertheless, the absolute value of this parameter was recoverable, i.e., the recovered parameters were also close to 1.

We also performed a model recovery test using the data simulated by the MB and MF algorithms. For each simulated dataset, we performed model comparison between the MB and MF algorithms based on the Bayesian Information Criterion (BIC). For all simulated participant datasets (bars in Fig. S1b), the “true” model that simulated the data was favoured (|Δ*BIC* |> 10), i.e., correctly recovered.

Finally, we performed an additional model recovery test regarding the hybrid-constant (HC) and hybrid-dynamic (HD) models. This specific test is performed to validate that, apart from differentiating MB behaviour from MF behaviour, whether our modelling approach can further identify the true trend of MF weights across trials at the group level. To this end, we ran autonomous simulations with the HC and HD algorithms separately for each participant using their best-fit parameters. Then, we fit both models (HC or HD) to the simulated data, and compared the models based on the overall BIC (sum of BICs across simulated participant datasets). For both the Visual-Haptic and Haptic conditions, the “true” model that simulated the data was favoured (|Δ*BIC*| > 10), i.e., the group-level trend of MF weight across trials was qualitatively recovered.

**Fig. S1.**
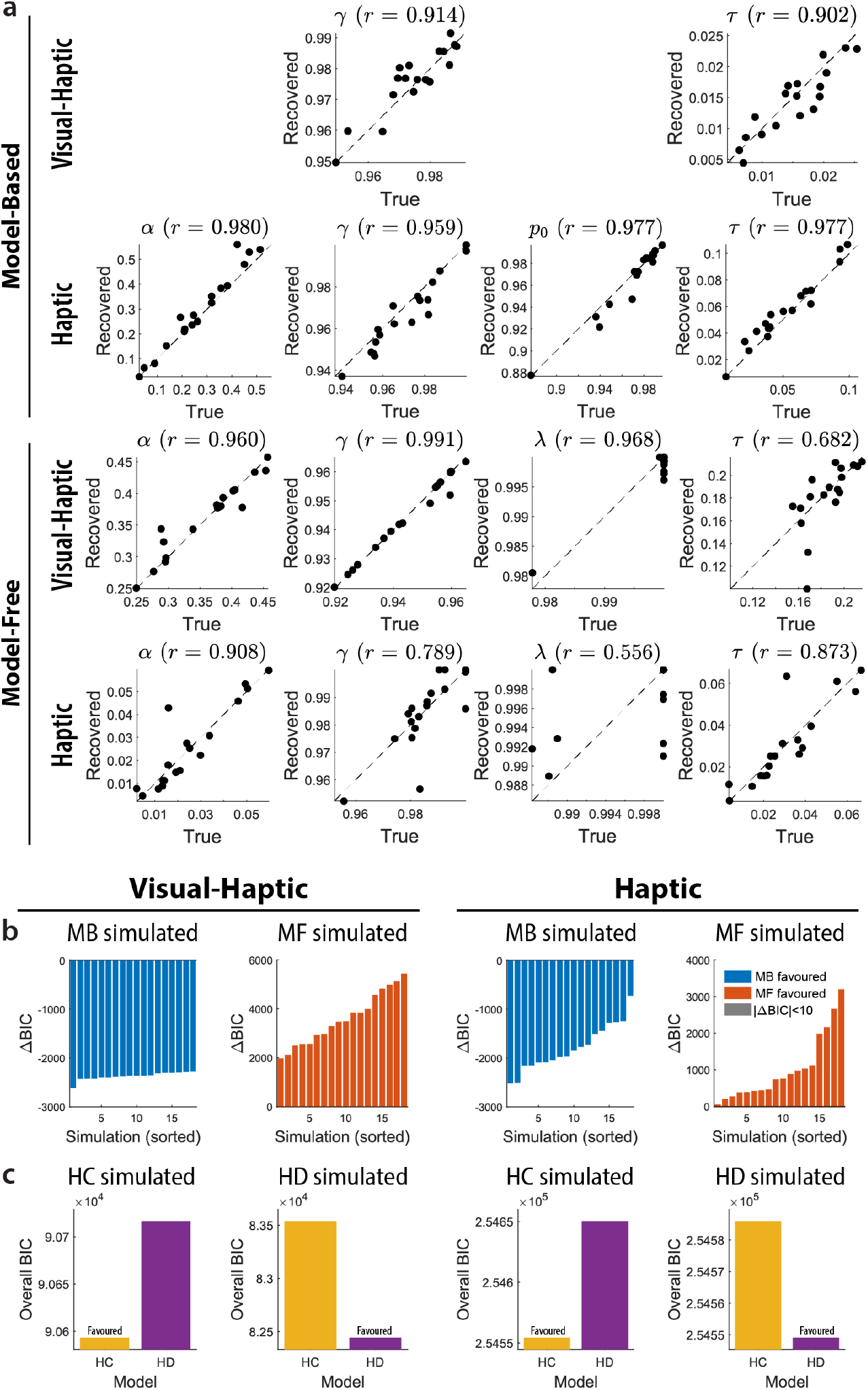
Model parameters and model identities are reliably recovered from simulated data. a, Parameter recovery for MB and MF algorithms in Visual-Haptic and Haptic conditions. Scatter plots show “recovered” vs. “true” parameters for each participant (dots) relative to the *x* = *y* unity line (dashed). b, Model recovery for MB and MF algorithms. Data simulated by each model were fit by both algorithms. The Δ*BIC* consistently favoured the “true” generating model (|Δ*BIC* |> 10) across all simulations. c, Group-level recovery for HC and HD models. Overall BIC (summed across participants) favoured the generating model in both experimental conditions. This confirms the modelling approach can correctly identify the trend of MF weights across trials.

### Hybrid action strategy divergence analysis

To confirm that the hybrid models meaningfully capture the relative contribution of MB and MF strategies, we compared the policies (probabilities of selecting each available action) generated by the MB and MF components in the hybrid models. The MF weights in the hybrid models are only interpretable when the MB and MF policies are generally distinguishable. Notably, because both components can degrade into a random strategy under specific parametrization (e.g., an extremely high Boltzmann exploration temperature parameter *τ*), we also compared both components against a random strategy.

We computed the Jensen–Shannon (JS) divergence between MB and MF policies, as well as between each policy and a uniform random policy, at each step. Figure S2 shows the JS divergence between each pair of strategies in each condition as a function of trial number, based on the hybrid-stepwise (HS) model. Results from the other two hybrid models, hybrid-constant (HC) and hybrid-dynamic (HD), are qualitatively similar.

In all three conditions (Visual-Haptic, Haptic, and the navigation task), the MB and MF strategies were consistently different from one another and from a random strategy (JS divergence above zero), for the majority of participants. However, for one participant in the Haptic condition, the MF component converged to a random policy (Fig. S2f). This specific participant exhibited the lowest overall MF weight in the HS model and the second-highest Bayesian evidence favouring an MB strategy over an MF strategy (based on Δ*BIC* between single MB and MF algorithms) in that condition. These metrics indicate that the hybrid model successfully identified the absence of MF behaviour in this participant; by effectively suppressing the MF weight, the model rendered the component’s internal parameters irrelevant to the likelihood function, leading to its “collapse” into a random strategy.

In sum, these results validate the meaningful interpretation of the MB and MF components and their respective weights within the hybrid modelling framework.

**Fig. S2.**
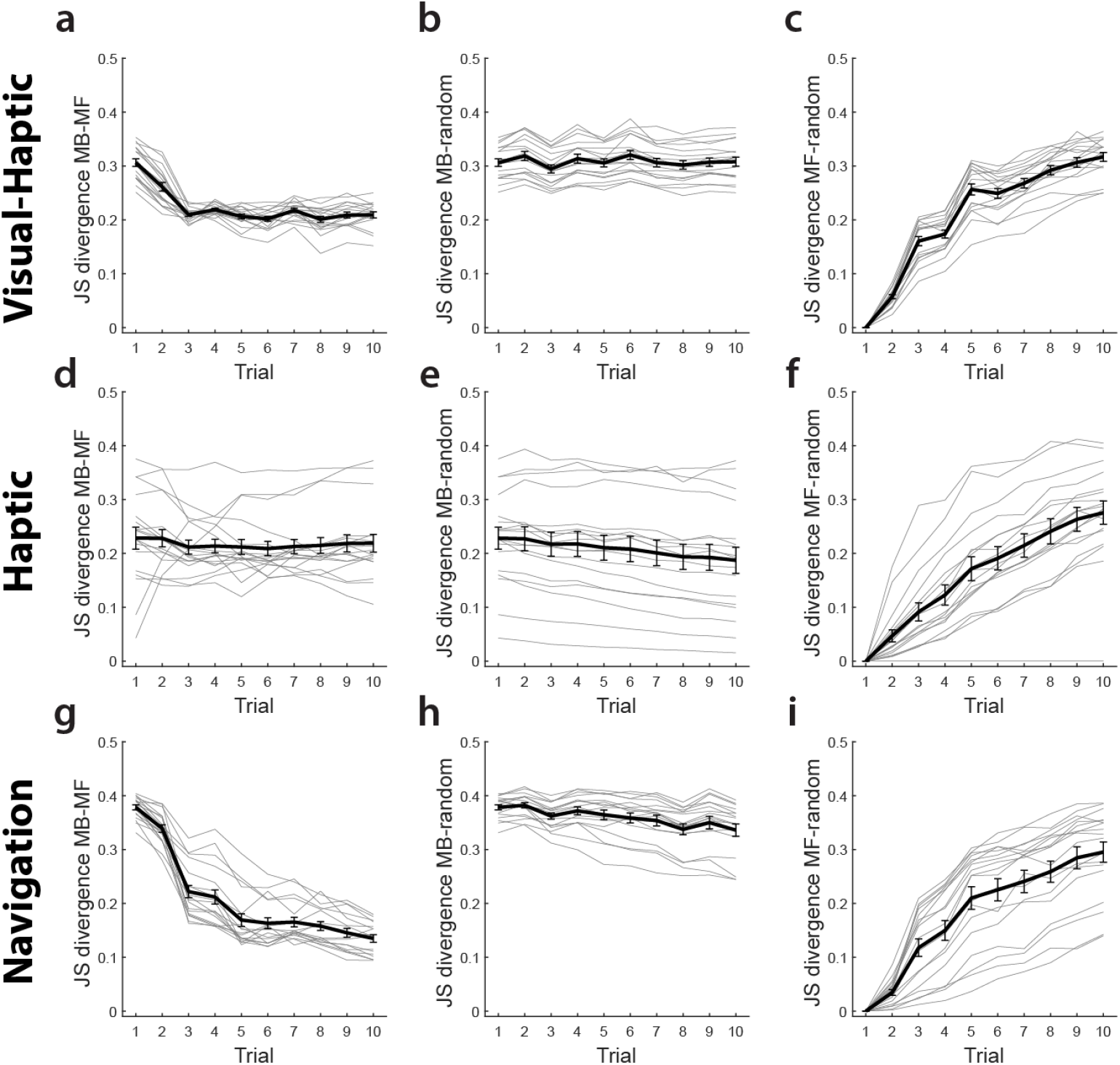
Model-based and model-free policies are consistently distinguishable within the hybrid framework. a–i, JS divergence across trials for the Visual-Haptic (a–c), Haptic (d–f) conditions in the reachable space task, and the navigation task (g–i). Comparisons include MB vs. MF policies (a, d, g), MB vs. random policies (b, e, h), and MF vs. random policies (c, f, i). Thick black lines represent the group mean, while thin grey lines represent individual participants. Error bars show *SEM*.

### Performance-optimized model-based and model-free algorithms

In Fig. 3m-p, we plotted performance of the MB and MF algorithms simulated using parameters fitted to human participants’ data. To evaluate the general performance of the two different algorithms in this maze task, without the constraint that they must produce human-like behaviour, we optimized each algorithm by searching for a set of parameters that minimized the average path length in the simulations. This analysis is based on the Haptic condition, where the maze layout is invisible.

Figure S3 shows the simulation results for the performance-optimized algorithms, together with human performance. The performance-optimized MB algorithm achieved a 100% success rate and an average path length close to the shortest path length possible under each maze configuration after approximately 2 trials of learning. This demonstrates that the optimized MB algorithm is highly effective at solving our maze learning task, even exceeding average human performance. In comparison, the performance-optimized MF algorithm improved across trials but still performed worse than MB or humans, highlighting the generally limited learning capacity of the MF algorithm despite parameter tuning.

**Fig. S3.**
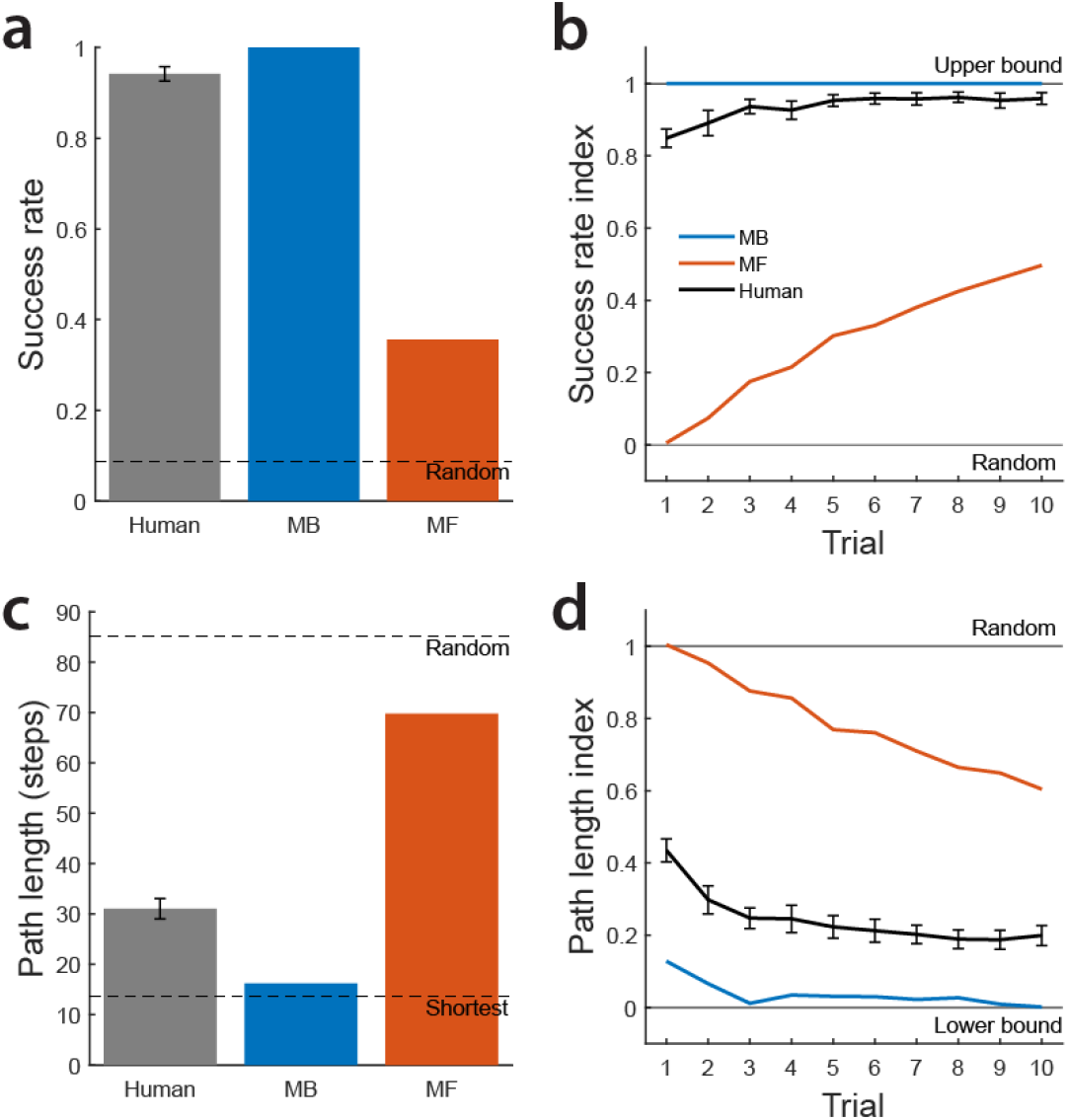
An optimized model-based algorithm matches or exceeds human performance whereas model-free alone remains limited. a. Overall success rate. b. Success rate index across trials, normalized between random performance (0) and the upper bound (1). c. Average path length in steps. The optimized MB algorithm achieves a path length near the shortest path, while MF remains much higher. d. Path length index across trials. Error bars show *SEM* across human participants.

### Successor representation algorithms

In our 10 × 10 grid maze environment, the successor representation (SR) is defined from a policy-dependent one-step transition matrix *T*_*π*_ and the successor matrix *M*. Importantly, *T*_*π*_ in the SR algorithm is distinct from the transition function *T* in our MB algorithm. In the MB algorithm, *T* (*s, a*) represents the probability, in the participant’s belief, of reaching the next state *s*^*′*^ (*s*^*′*^ is not blocked) after taking action *a* from state *s*, reflecting a mental map of the environment. In the SR algorithm, by contrast, *T*_*π*_(*s, s*^*′*^) gives the probability of transitioning to state *s*^*′*^ in one step from state *s* under policy *π*, reflecting the participant’s strategy for choosing actions.

The successor matrix *M* encodes the expected temporally discounted future occupancies under policy *π*:

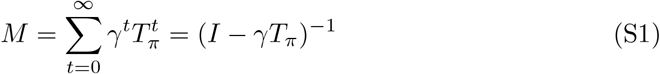

where *t* is the number of steps into the future, *γ* is the temporal discount factor, and *I* is the identity matrix (with ones on the diagonal and zeros elsewhere).

For each new maze, the initial one-step transition matrix *T*_*π*0_ is defined as a uniform random walk over the four possible actions (north, east, south, or west), assuming no maze blocks. For states at edges or corners, probabilities are evenly distributed among valid (i.e., within boundary) neighbours. To account for the assumption that one of the 100 states is absorbing—although its location is unknown at initialization—each row of *T*_*π*0_ was scaled to sum to 0.99:

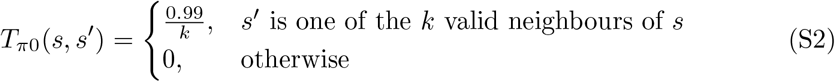

where *k ∈ {*2, 3, 4*}* is the number of valid neighbours. The initial successor matrix *M*_0_ is then calculated from *T*_*π*0_:

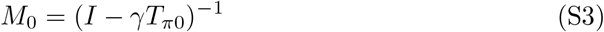

In the SR-TD algorithm, the successor matrix *M* is updated at each step using TD learning with eligibility traces. Specifically, the eligibility trace is initialized over all states at the beginning of each trial:

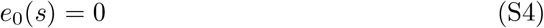

and updated at each step *t*:

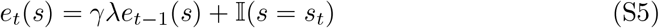

where *λ* is the memory decay factor and 𝕀 is the indicator function, which takes the value of 1 if the condition holds, and 0 otherwise. The successor matrix *M* is updated as:

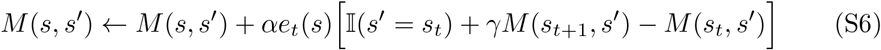

where *α* is the learning rate.

The SR algorithm separately encodes the target location with a reward function *R*:

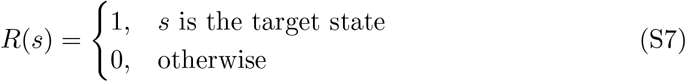

Given the successor matrix *M* and the reward function *R*, the value function *V* for each state *s* is computed as the expected discounted sum of future rewards:

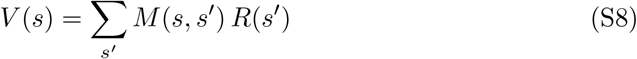

During action selection, the algorithm makes decisions based on the value function *V*. The probability of selecting each action option *a*_*i*_ from the current state is:

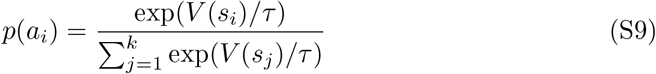

where *s*_*i*_ is the intended next state by taking action *a*_*i*_, *k* is the number of action options, and *τ* is the temperature parameter for Boltzmann exploration.

In the SR-MB algorithm, for each new maze, the one-step transition matrix *T*_*π*_ is initialized the same way as in the SR-TD algorithm. In contrast, at each step *t*, the SR-MB algorithm updates *T*_*π*_ based on the participant’s action taken, rather than directly updating the successor matrix *M* :

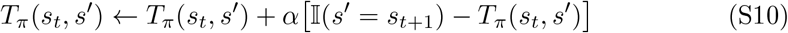

Thus, *T*_*π*_ estimates the participant’s action policy by tracking the historical frequencies of chosen next states.

After updating *T*_*π*_, the successor matrix is recomputed at each step:

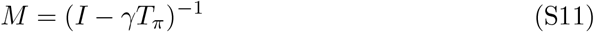

This enables *M* to be globally updated from *T*_*π*_, implementing MB planning rather than incremental TD updates as in SR-TD.

After computing the successor matrix *M*, the SR-MB algorithm combines it with the reward function *R* to obtain the value function *V* and selects actions based on *V* using the same procedure as in the SR-TD algorithm.

The SR-TD algorithm has 4 free parameters: *α, γ, λ*, and *τ*. The SR-MB algorithm has 3 free parameters: *α, γ*, and *τ*. We fit the two algorithms separately for each participant using maximum likelihood estimation (MLE).

Both the SR-TD and SR-MB algorithms we implemented are identical in the Visual-Haptic and Haptic conditions. Specifically, these SR algorithms are strictly policy-based, rather than environment-based as in the pure MB algorithm. Therefore, the SR algorithms can learn only from action experience and cannot use the visual information of the maze layout provided in the Visual-Haptic condition, unlike the MB algorithm.

Fig. S4 compares, in the Visual-Haptic and Haptic conditions, the two SR algorithms with the MB and MF algorithms as well as the three hybrid models (HC, HD and HS) based on the Fixed-Target condition.

Fig. S4a and c show the overall BIC (sum of BICs across all participants) for each model. We did not calculate BIC for the HS model due to its non-parametric nature regarding the MF weights. For both conditions, the SR-TD algorithm has the highest BIC (i.e., worst) among all models, while the SR-MB lies between the MB and MF algorithms. Neither of the SR models is favoured over the two hybrid models, HC and HD. Fig. S4b and d show the likelihood per step as a function of trials for each algorithm (averaged across different mazes and participants). Both SR algorithms have lower likelihood than the three hybrid models across all 10 trials.

Note that the SR algorithms can be viewed as representation-level hybrid models combining features of MB and MF learning. Our results suggest that such representation-level hybrid models are less capable than our mixture-of-expert hybrid models in terms of explaining participants’ behaviour in our tasks.

**Fig. S4.**
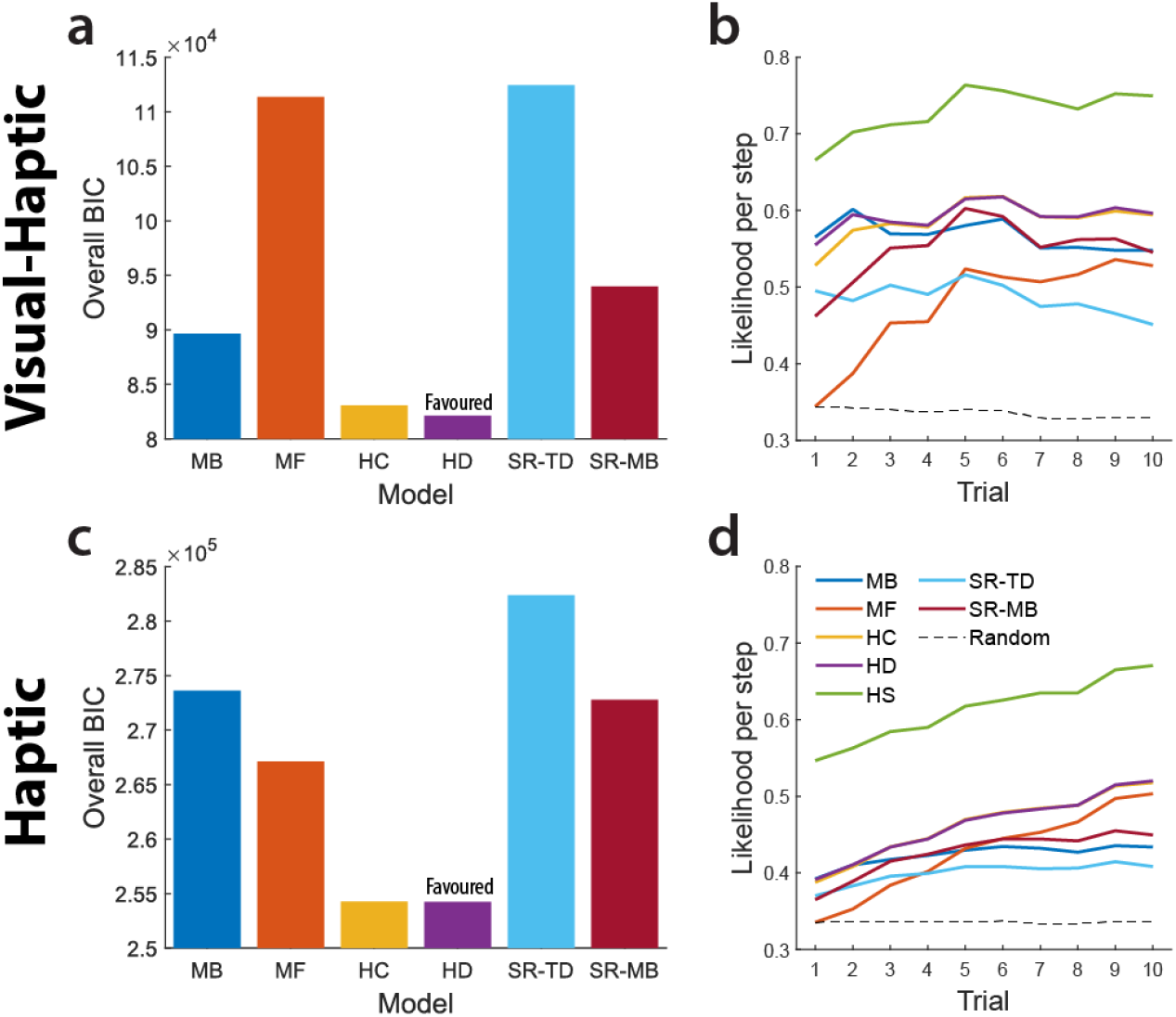
Successor representation algorithms are outperformed by hybrid models in the reachable space task. a, c, Overall BIC (sum across participants) for the Visual-Haptic and Haptic conditions. The SR-TD algorithm shows the highest (worst) BIC, while the SR-MB model performs between the MB and MF algorithms. Neither SR-based model is favoured over the hybrid-constant (HC) or hybrid-dynamic (HD) models. b, d, Likelihood per step across trials for each algorithm in the Visual-Haptic and Haptic conditions. Both SR models consistently exhibit lower likelihood compared to the three hybrid models (HC, HD, and HS) across all trials.

### Potential effect of sex on action strategies

Previous studies have reported sex-related differences in human spatial cognition and navigation strategies. Considering this, we tried to balance the number of female and male participants in our study (Visual-Haptic: 11 females, 7 males; Haptic: 9 females, 9 males). We also performed a two-way ANOVA for unbalanced design to assess the effect of condition (Visual-Haptic vs. Haptic) while accounting for the potential effect of sex on the overall MF weight.

We found a significant main effect of condition (*F* (1, 32) = 15.39, *p* = 10^*−*4^), demonstrating that the overall MF weight is significantly higher in the Haptic condition than in the Visual-Haptic condition. Meanwhile, the main effect of sex did not reach the significance level (*F* (1, 32) = 3.57, *p* = 0.068). Moreover, the interaction between condition and sex was non-significant (*F* (1, 32) = 2.70, *p* = 0.11), which statistically confirms that the effect of condition is consistent for both female and male participants. These findings indicate that the observed action strategy difference is a direct result of the condition and is independent of participant sex.

**Fig. S5.**
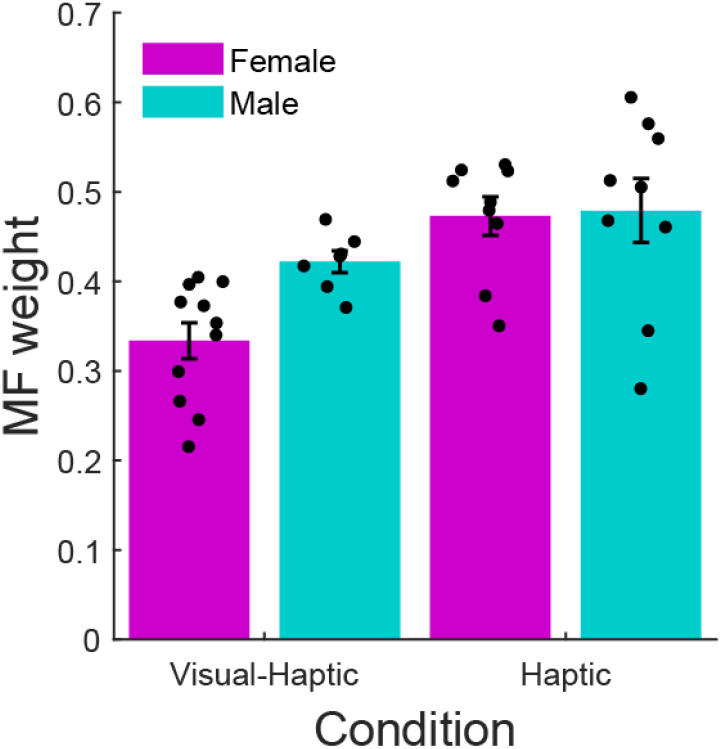
The condition difference in model-free reliance is independent of participant sex. Bars show overall MF weight, averaged across participants, categorized by sex for the Visual-Haptic and Haptic conditions. Dots represent individual participants, and error bars show *SEM*. While the Haptic condition shows a significantly higher MF weight than the Visual-Haptic condition, the main effect of sex and its interaction with condition were statistically non-significant.

## Notes

### Competing Interest Statement

The authors have declared no competing interest.

### Funding

- Natural Sciences and Engineering Research Council of Canada: RGPIN/05944-2019
- Canadian Institutes of Health Research: 156173
- National Institutes of Health: R01NS117699
- Air Force Office of Scientific Research: FA9550-22-1-0337

